# Transcriptional regulators predicted to drive macrophage dysregulation during impaired wound healing in diabetic mice

**DOI:** 10.64898/2026.04.21.719960

**Authors:** Brandon Lukas, Jingbo Pang, Yang Dai, Timothy J. Koh

## Abstract

Dysregulation of Mo/M*φ* activity is known to contribute to impaired healing in diabetes; however, the mechanisms underlying this dysregulation are not well understood. In this study, we used a variety of bioinformatics approaches along with our time series scRNA-seq data on wound Mo/M*φ* from non-diabetic and diabetic mice to identify transcriptional regulators (TRs) that drive Mo/M*φ* state transitions during normal and impaired healing. First, we used the Lamian framework and our newly developed Pseudotime Graph Diffusion method to show that state transitions from early stage phenotypes to later stage reparative and antigen presenting phenotypes characteristic of normally healing wounds are impaired and that transitions to inflammatory, foam cell-like, and Lyve-1^+^ M*φ* phenotypes are enhanced during impaired healing of diabetic mice. Using our BITFAM model, we identified a broad range of TRs predicted to be preferentially active in each cell state and using CellOracle, we performed in silico perturbation to identify groups of TRs predicted to drive cell state transitions along multiple trajectories (e.g. CEBPA, IRF8), whereas other TRs were predicted to drive cell state transition towards reparative phenotypes (e.g. NR1H3, NR3C1) or towards an antigen-presenting phenotype (e.g. IRF4, OGT). Selected findings were validated using existing experimental data, confirming the usefulness of this approach. In conclusion, we identified TRs that likely drive Mo/M*φ* state transitions towards desirable and undesirable phenotypes for wound healing. These findings provide insight into novel targets for altering Mo/M*φ* phenotypes to promote healing of diabetic wounds.

## 1 Introduction

Chronic wounds associated with diabetes are an escalating problem around the world [1]. People suffering from diabetes incur a 25% lifetime risk of developing chronic wounds, and associated risk of amputation, decreased quality of life, high morbidity and mortality [2–4]. Effective wound healing requires coordinated responses of different cell types during overlapping phases of inflammation, proliferation and remodeling [5, 6] and chronic wounds are known to exhibit defects in each phase of healing, resulting in a persistent inflammatory state [7, 8]. Monocytes and macrophages (Mo/M*φ*) play essential roles in wound healing [9, 10], and dysregulation of Mo/M*φ* activity contributes to impaired healing in diabetes [11, 12]. However, the mechanisms underlying this dysregulation are not well understood.

Over the course of wound healing, Mo/M*φ* adopt a wide spectrum of functional phenotypes that is best characterized by the resolution afforded by single cell RNA-seq (scRNA-seq) [13–16]. Transcription factors (TFs) and other transcriptional regulators (TRs), including TF modifiers and epigenetic regulators, play critical roles in regulating gene expression involved in specifying this spectrum of Mo/M*φ* phenotypes [17–20]. We recently reported that, over the course of normal wound healing, wound Mo/M*φ* cluster into early-stage Mo/M*φ*, late-stage M*φ* and antigen presenting cells [16] and that each cluster showed differential chromatin accessibility and differential TR activity predicted by our BITFAM model [21, 22]. Network analysis revealed highly connected TR communities that cooperate to regulate wound Mo/M*φ* phenotypes. Importantly, we validated a pro-inflammatory role for NR4A1 during wound healing, showing that *Nr4a1* knockout mice exhibit decreased inflammatory gene expression in early-stage wound Mo/M*φ*, along with delayed wound re-epithelialization and impaired granulation tissue formation. Although a number of reports have characterized dysregulation of wound Mo/M*φ* phenotypes during impaired diabetic wound healing, the role of altered TR activity in such dysregulation has not been elucidated.

In the present study, we used the Lamian framework [23] and our newly developed Pseudotime Graph Diffusion method on our time-series scRNA-seq data from skin wound cells of non-diabetic and diabetic mice to show that wound Mo/M*φ* exhibit altered state transitions from early to later stage phenotypes in diabetic versus non-diabetic mice, including impaired transitions to reparative and antigen-presenting phenotypes and enhanced transitions to inflammatory, foam cell-like, and Lyve-1^+^ phenotypes. Using our BITFAM model [21, 22], we identified a broad range of TRs that are predicted to be preferentially active in each cell state, and those that are dynamically active over specific ranges of pseudotime. Using CellOracle [24], we performed in silico TR perturbation to predict which TRs drive the various cell state transitions in non-diabetic and diabetic wounds. Finally, we validated selected findings using existing experimental data, confirming the usefulness of this approach to identify TRs that drive state transitions in diabetic wounds.

## 2 Methods and Materials

### Animals

C57BL6/J (Non-diabetic, ND) and BKS.Cg-*Dock7* ^m +/+^ *Lepr* ^db^/J (a model of type 2 diabetes, DB) were purchased from the Jackson Laboratory (Bar Harbor, ME) and housed for at least two weeks before experiments. All animal procedures were approved by the Animal Care and Use Committee of the University of Illinois Chicago.

### Wound model

Adult male mice (age 10-12 weeks) were subjected to excisional skin wounding with an 8 mm biopsy punch as described [16, 25]. Skin wounds were collected using a 12 mm biopsy punch, then dissociated by mechanical and enzymatic digestion into single cell suspension as previously described [16, 25].

### scRNA-seq

Wound cells were pooled from 4 wounds of male ND or DB mice each on day 3, 6, and 10 post-injury. Sorted live CD11b^+^CD45^+^Ly6G^*−*^cells were processed for scRNA-seq using the 10x Chromium Next GEM Single Cell 3’ Reagent Kits (10X Genomics, San Francisco, CA, USA). Libraries were sequenced on HiSeq Sequencing Systems (Illumina, San Diego, CA, USA) with paired-end reads aiming for 100,000 reads/cell [25].

After demultiplexing, cells with *>* 10% mitochondrial gene expression, unique features/gene expression *<* 500 and *>* 10,000, or *<* 1,000 UMI counts were omitted and final 6,109 cells (ND-D3: 961, ND-D6: 1,121, ND-D10: 1,214, DB-D3: 1,334, DB-D6: 894, and DB-D10: 585) were included for downstream analysis [25]. scRNA-seq data used in this study are available in Gene Expression Omnibus series record GSE203244 and GSE302547.

### scRNA-seq data processing

Raw count matrices were processed using Seurat v5 [26] with default parameters unless otherwise noted. Expression data were log-normalized, and the top 2,000 highly variable genes (HVGs) were selected. UMAP embeddings were computed from the first 30 principal components. Cluster-specific marker genes were identified using a one-versus-rest framework, and gene set enrichment analysis (GSEA) was performed against the MSigDB Hallmark collection using GSEApy [27] with preranked analysis (1,000 permutations), ranking genes by sign(log_2_ fold change) *× −*log_10_(adjusted *P*-value).

### Trajectory inference and pseudotime analysis

Lamian [23] was used for trajectory inference and differential pseudotime analysis. Cells were clustered in PCA space via *k*-means (*k* = 12, selected by elbow method), and a minimum spanning tree was constructed over cluster centroids, rooted at day 3 cells. Branch robustness was assessed by permutation (1,000 permutations). Pseudotime-dependent gene expression was identified using Lamian’s time-dependent expression (TDE) test, applied branch-wise to the top 5,000 HVGs per branch. Significant genes (FDR *<* 0.05) were classified as early, middle, or late based on peak expression timing (top 1%, middle, or bottom 1% of branch pseudotime, respectively) and subjected to pathway enrichment against the MSigDB Hallmark collection.

### Enhanced trajectory visualization

Low-dimensional embeddings such as UMAP may not faithfully preserve trajectory structures supported in higher-dimensional PCA space. To address this issue, we applied Pseudotime Graph Diffusion (PGD) [28], a method we developed that smooths cell-level embeddings along a pseudotime graph via Markovian diffusion. Seurat PCA embeddings were diffused along the Lamian-inferred pseudotime graph (*α* = 0.6, with self-loops) and subsequently projected to UMAP for improved trajectory visualization.

### Transcriptional regulator activity inference

TR activities were inferred using BITFAM [22], a Bayesian factor analysis model that incorporates a gene regulatory network (GRN) as prior knowledge. The GRN used in this study was derived from ChIP-Atlas [29] TR–target gene associations (±5 kb; experiments annotated as “dendritic”, “macrophage”, or “monocyte”; downloaded May 14, 2025). Target genes of Zbtb46 were manually added given its established role in dendritic cell differentiation [30]. The GRN was pruned to exclude TRs detected in fewer than 100 cells (-3 TRs) or with fewer than 10 target genes (-6 TRs) among the top 2,000 HVGs. BITFAM was run four times for robustness; nominal *P*-values from one-versus-rest comparisons were combined across runs using a signed Stouffer *Z* -score method with Benjamini–Hochberg FDR correction. Per-cell TR activity was summarized as the median across runs.

### In silico transcriptional regulator perturbation

CellOracle [24] was used to simulate TR knockout (expression set to zero) and overexpression (expression set to the safe-range upper bound, calculated as the observed maximum plus the full expression range (max + (max −min)) of single-cell expression values for each gene). Perturbation scores, computed as the inner product of simulated perturbation vectors with pseudotime-derived developmental vectors, were summarized per branch by separately summing positive (promoting transitions) and negative (inhibiting transitions) scores. Markov random walk simulations (500 steps) were additionally performed to quantify perturbation-induced changes in cluster occupancy, reported as percent change relative to initial walker counts

## 3 Results

### Mo/M*φ* cell states are altered in diabetic wounds

We recently reported that networks of TR guide wound Mo/M*φ* through state transitions from pro-inflammatory to pro-healing phenotypes as normal healing progresses in ND mice [16]. In the present study, we used the Lamian computational framework [23] to identify dysregulation of state transitions of wound Mo/M*φ* in DB mice (Fig. 1). We also developed and applied Pseudotime Graph Diffusion (PGD), a post hoc technique that enhances the visualization of pseudotemporal trajectory structure in low-dimensional representations through graph-based diffusion without altering the underlying data, facilitating downstream analyses. Next, we used our BITFAM model to predict TR activity that may contribute to dysregulation of state transitions [21, 22]. Finally, we used CellOracle to identify the impact of in silico perturbation of TR expression and validated the predicted effects of TR knockout using prior data from our laboratory and others [24].

**Figure 1.**
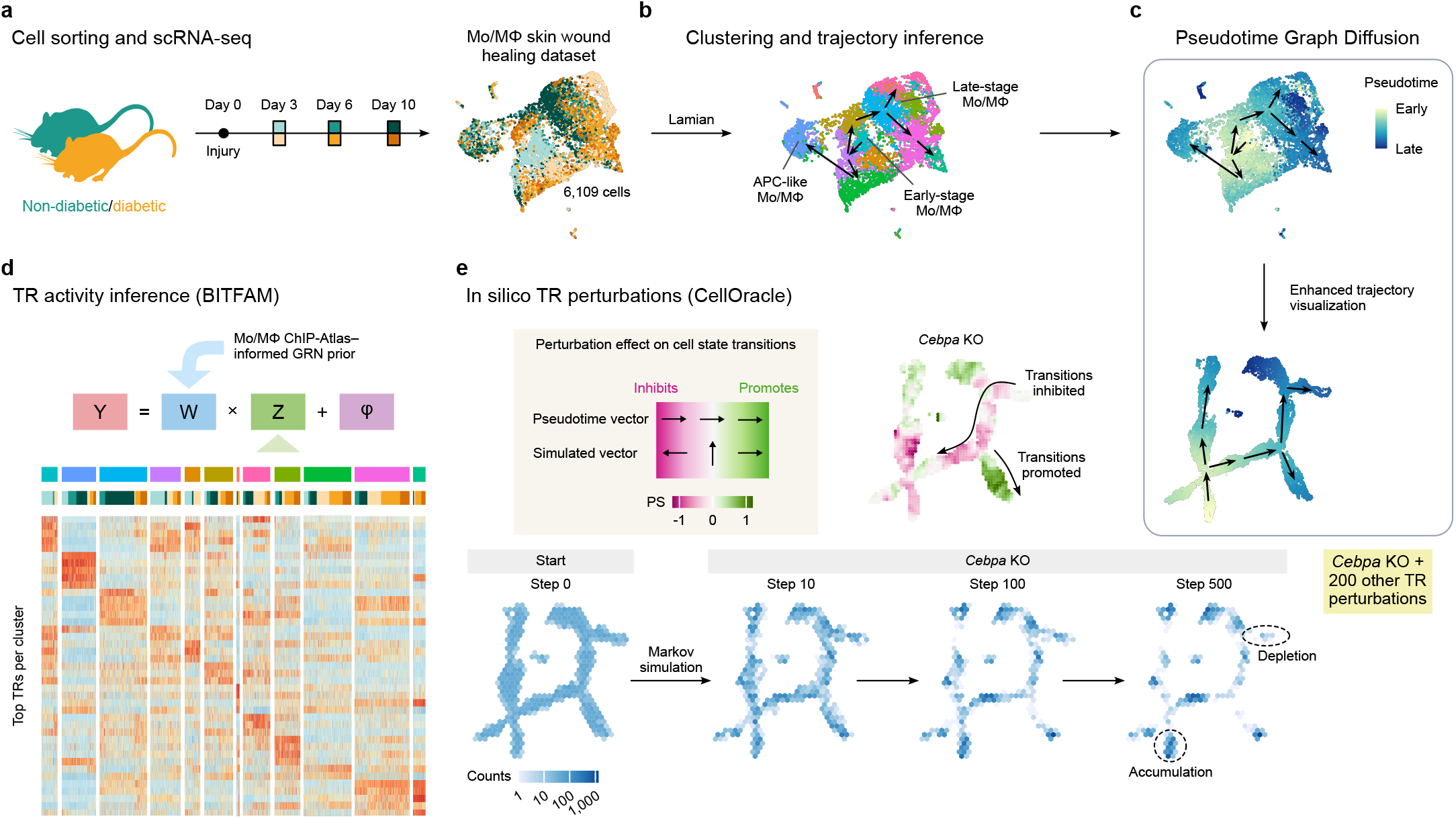
Experimental design and bioinformatics analysis workflow. **a**. Schematic representation of experimental design. Non-diabetic and diabetic mice were injured, monocyte/macrophage populations were isolated from the wound on days 3, 6, and 10 post-injury and subjected to scRNA-seq using the 10x Chromium platform. **b**. Clustering and trajectory inference was performed to identify cell states and model state transitions using the Lamian framework [23]. **c**. Our new pseudotime graph diffusion (PGD) method was applied to enhance the inferred trajectory visualization. **d**. Activity of transcriptional regulators was inferred using a Bayesian inference model (BITFAM [22]) integrating a custom monocyte/macrophage ChIP-Atlas–informed prior network. **e**. Systematic in silico TR perturbations for 98 TRs were performed with CellOracle [24], including both knockout (KO) and overexpression simulations, to predict consequences on cell state transitions through perturbation score (PS) analyses and Markov simulations. Illustration from NIAID NIH BIOART Source bioart.niaid.nih.gov/bioart/372.

First, clustering of pooled scRNA-seq data from live CD45^+^CD11b^+^Ly6G^*−*^cells isolated from ND and DB skin wounds on days 3, 6 and 10 post-injury identified 12 clusters of cells (Fig. 2a,b; cluster numbers arbitrarily assigned). Clusters 12, 5, and 3 were populated preferentially by cells from ND wounds (Fig. 2c): cluster 12, populated primarily by ND day 3 cells, expressed an interferon activation signature (*e.g. Rsad2, Slfn4, Ifi205*; Fig. 2d shows the top 5 marker genes, see Supplementary Data S1 for the full set of differentially expressed genes). Cluster 5, populated primarily by ND day 6 and 10 cells, expressed an antigen presenting cell (APC) signature (*e.g. Cd209a, Ciita, MHCII* genes). Cluster 3, populated primarily by ND day 10 cells, expressed a mixed reparative signature (*e.g. Arg1, Vegfa, Chil3*). In contrast, clusters 4, 7, 1 and 8 were populated preferentially by cells from DB wounds (Fig. 2c): Clusters 4 and 7, composed primarily by DB days 3 and 6 cells, expressed mixed complement-associated (*e.g. C1qa,b,c, Mrc1, Il1b*) and inflammatory (*e.g. Cxcl2, Tnfaip, Ptgs2*) phenotypes, respectively. Cluster 1, populated preferentially by DB day 6 and 10 cells, expressed a lipid handling, foam cell-like phenotype (*e.g. Lpl, Lipa, Cd36*). Cluster 8, populated primarily by DB day 6 and 10 cells, expressed a Lyve-1 M*φ* signature (*e.g. Lyve1, Folr2, Cd163*).

**Figure 2.**
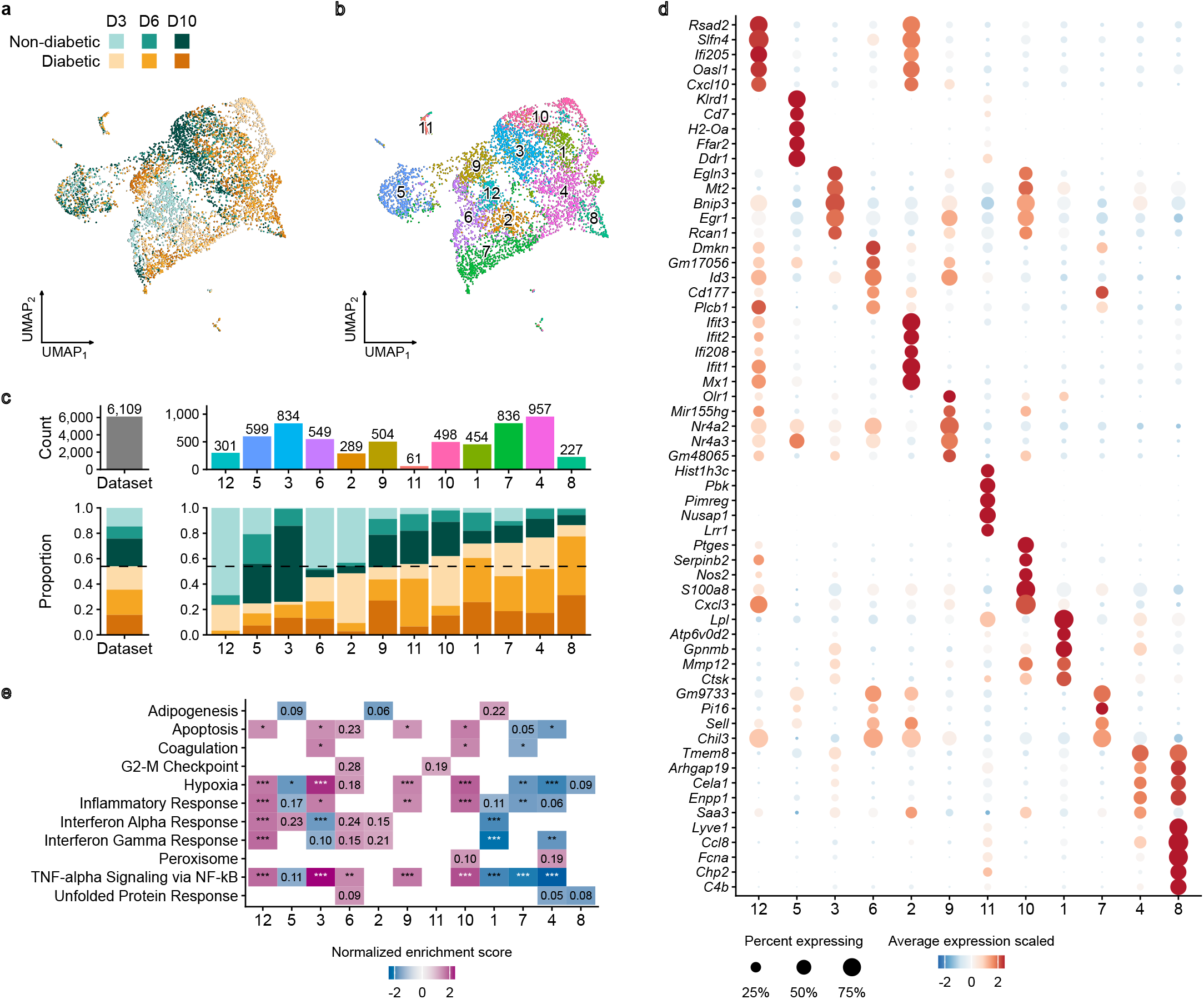
Monocyte/macrophage cell states are dysregulated in wounds of diabetic mice in all phases of healing. **a**. UMAP visualization of cells colored by sample, days 3 (D3), 6 (D6) and 10 (D10) post-injury for non-diabetic (ND) and diabetic (DB) mice. **b**. UMAP visualization of cells colored by cluster (clusters 1-12). **c**. Cell counts (top) and proportions of each sample (bottom) that contribute to each cluster. **d**. Dot plot visualization of top 5 differentially expressed genes ranked by average log2FC per cluster in one-vs-rest analysis (note that the full list of differentially expressed genes can be found in Supplementary Data S1). **e**. Heatmap visualization of normalized enrichment scores of selected MSigDB Hallmark 2020 terms from gene set enrichment analysis (GSEA). Shown are scores with *P <* 0.05; values denote FDR *Q*-values (\**Q <* 0.05, \*\**Q <* 0.01, \*\*\**Q <* 0.001).

Gene set enrichment analysis confirmed characterization of the cluster phenotypes discussed above and provided additional insight (Fig. 2e; see Supplementary Data S2 for full analysis, including lead genes for each pathway). Clusters 12, 6 and 2, all populated primarily with day 3 cells from both ND and DB mice, showed preferential activity of interferon and tumor necrosis factor (TNF)/nuclear factor (NF)-*κ*B pathways, although these were only trends in clusters 6 and 2. Cluster 1, populated primarily with DB day 3 and 6 cells, showed preferential activity of the adipogenesis pathway. Clusters 4 and 7, populated primarily with DB cells from all time points showed preferentially reduced activity of the apoptosis pathway, consistent with our previous observation that reduced apoptosis of DB wound Mo/M*φ* contributes to their persistence in wounds of diabetic mice [31]. Clusters 4 and 10, also populated primarily with DB cells from all time points, showed trends of preferential activity of the peroxisome pathway, perhaps indicating an adaptive response to oxidative stress. Finally, clusters 1, 4 and 7, again populated primarily with DB cells from all time points, showed preferentially low activity of interferon, TNF, inflammatory and hypoxia pathways consistent with previous finds of dysregulation of inflammatory pathways in human DB wound Mo/M*φ* [32].

### Pseudotime analysis reveals inhibited/altered Mo/M*φ* cell state transitions in diabetic wounds

Next, we used Lamian [23] to infer a pseudotemporal trajectory followed by PGD to enhance its visualization. Two trajectories were identified with high confidence (Fig. 3a,b): the first branch started in cluster 12, then transitioned through clusters 6 and 7, and terminated in cluster 5 (branch 1: 12-6-7-5). The second branch started in cluster 12, and transitioned through clusters 6 and 9 before terminating in clusters 3 and 10 (branch 2: 12-6-9-3-10). A lower confidence branch connected cluster 12 to clusters 6, 9 and 3 and then to clusters 4 and 8 (branch 3: 12-6-9-3-4-8).

**Figure 3.**
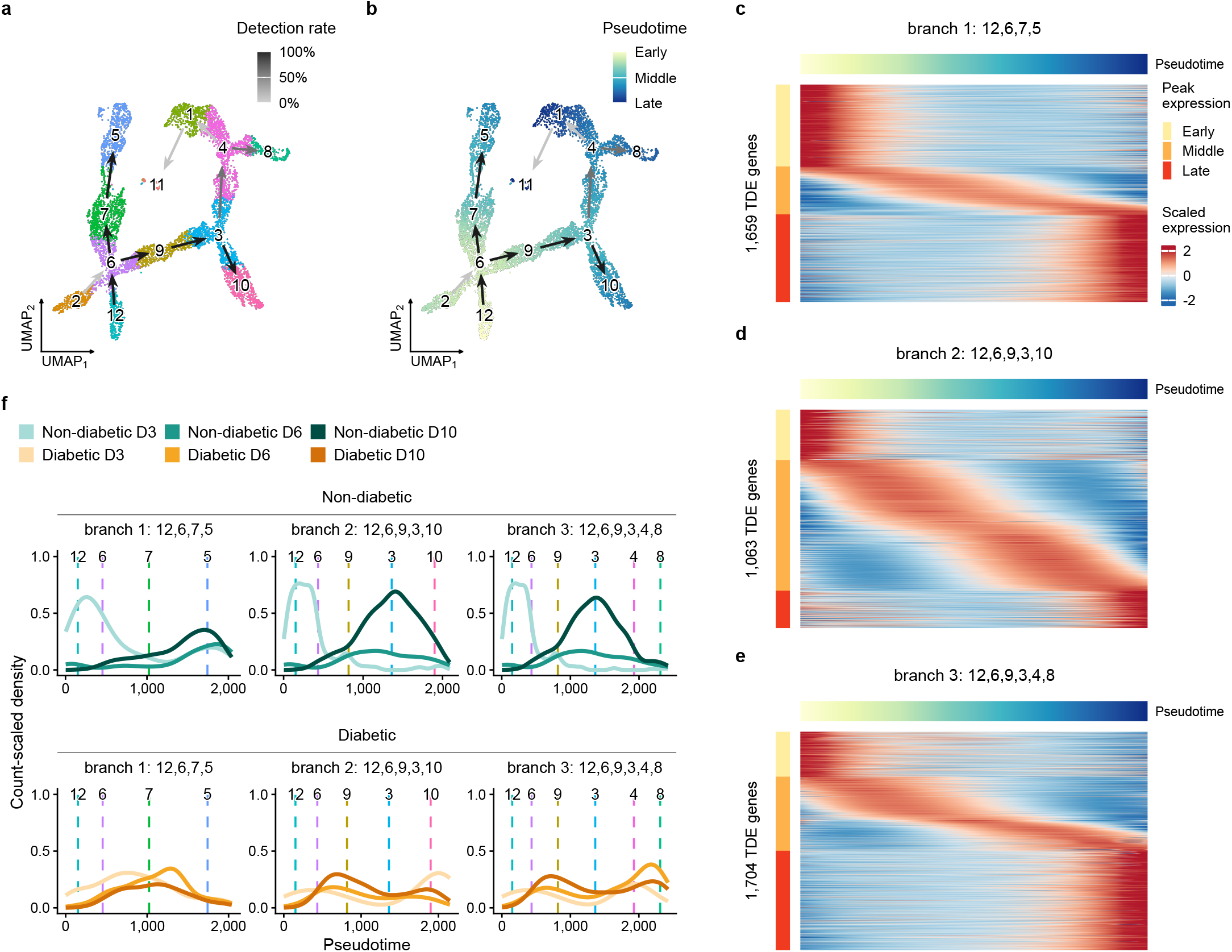
Trajectory inference suggests delayed or altered cell state transitions in diabetic wound monocytes/-macrophages. Trajectory inference performed using the Lamian framework followed by enhanced trajectory visualization via PGD smoothing. **a**. UMAP visualization of cells colored by cluster with cluster-based pseudotemporal branches. Branch lines colored by detection rate from bootstrap resampling; dark lines represent trajectories detected with high confidence. Lighter lines represent trajectories detected with lower confidence. **b**. UMAP visualization of cells colored by pseudotime; low pseudotime values correspond to early wound cell states, higher pseudotimes correspond to different cell differentiation trajectories. **c**. Density of non-diabetic and diabetic cells along pseudotime of branch 1: 12,6,7,5 and branch 2: 12,6,9,3,10. Dashed lines indicate median pseudotime of clusters along branches. **d**. Heatmap visualization of temporally differentially expressed (TDE) genes along pseudotime for branch 1: 12,6,7,5. **e**. Heatmap visualization of TDE genes along pseudotime for branch 2: 12,6,9,3,10. **f**. Heatmap visualization of TDE genes along pseudotime for branch 3: 12,6,9,3,4,8.

Each branch showed dynamic expression of genes with subsets showing preferential expression during early, middle and late pseudotimes (Fig. 3c,d,e see Supplementary Data S3 for full list of dynamically expressed genes along each branch). Gene set enrichment analysis showed preferential enrichment of the interferon *α* and *γ* pathways at early pseudotimes for all trajectory branches (Fig S1). Branch 1 showed 1,659 dynamically expressed genes (FDR *<* 0.05), with a relatively rapid switch from early to late phenotypes over pseudotime, and preferential enrichment of the IL2/STAT5 pathway at late pseudotimes, perhaps indicating the importance of STAT5 in state transitions or functions of APC phenotype cells in wounds [33, 34]. Branch 2 showed 1,063 dynamically expressed genes, with a more gradual switch from early to late phenotypes. Branch 2 showed preferential enrichment of epithelial-mesenchymal transition and angiogenesis at intermediate pseudotimes, reflecting the reparative phenotype of cells that populated these pseudotimes, but no pathways were significantly enriched at late pseudotimes, perhaps reflecting the mixed phenotype of the DB cells that populated late pseudotimes. Finally, branch 3, identified with less confidence, showed 1,704 dynamically expressed genes with a large number of these expressed at late pseudotimes. Branch 3 showed preferential enrichment of adipogenesis, fatty acid metabolism, and xenobiotic metabolism pathways at late pseudotimes, highlighting the altered metabolism of the DB cells that populated late pseudotimes of this branch.

When comparing the density of Mo/M*φ* cells along the trajectory branch 1 (ending in cluster 5), Mo/M*φ* from ND wounds were concentrated in the early and late portions of the trajectory with a terminus in cluster 5, which had an APC phenotype, whereas cells from DB wounds were concentrated in the middle of the trajectory (Fig. 3f). These results may reflect both a reduced number of cells with the early inflammatory phenotype and an incomplete transition to the APC phenotype in DB wounds, with cells “stuck” in cluster 7, which exhibited a mixed inflammatory phenotype. When comparing the density of Mo/M*φ* cells along the trajectory branch 2 (ending in clusters 3 and 10), Mo/M*φ* from ND wounds were concentrated in early and middle portions of the trajectory, with an apparent terminus in cluster 3, which had a reparative phenotype, whereas cells from DB wounds were concentrated at shifted pseudotimes, indicating altered transitions in DB wounds (Fig. 3f). Interestingly, cluster 10 appeared to be a terminus of trajectory branch 2, but was populated primarily by day 3 cells from DB wounds, and had a mixed inflammatory/reparative phenotype, expressing high levels of *Nos2, Tnf* and *Arg1* (Fig. S1). This may reflect dysregulation of early wound Mo/M*φ* in DB mice, with impaired inflammatory function, as has been found in human diabetic foot ulcers [32]. Branch 3 also showed dysregulation of DB wound Mo/M*φ* at both early and late pseudotimes, with increased transitions to cluster 8 in DB day 6 and 10 cells, which expressed high levels of *Il1b, Mrc1* and *Igf1* along with Lyve-1 M*φ* markers (Fig. 3f, S2); cells with this phenotype were rare in ND wounds. In short, this analysis indicates dysregulation of DB wound throughout the time course analyzed.

### TRs show differential inferred activity between Mo/M*φ* cell states

Next, we used our BITFAM model to infer the activity of TRs that potentially drive the observed dysregulation in Mo/M*φ* cell states in diabetic wounds. Distinct groups of TRs showed differential inferred activity in different Mo/M*φ* clusters (Fig. 4a, Supplementary Data S4). Interestingly, levels of inferred TR activity generally did not correlate well with levels of mRNA expression (Fig. 4b, Supplementary Data S5). These findings corroborate those our previous study [21], that mRNA expression is necessary but not sufficient for high inferred TR activity predicted by BITFAM.

**Figure 4.**
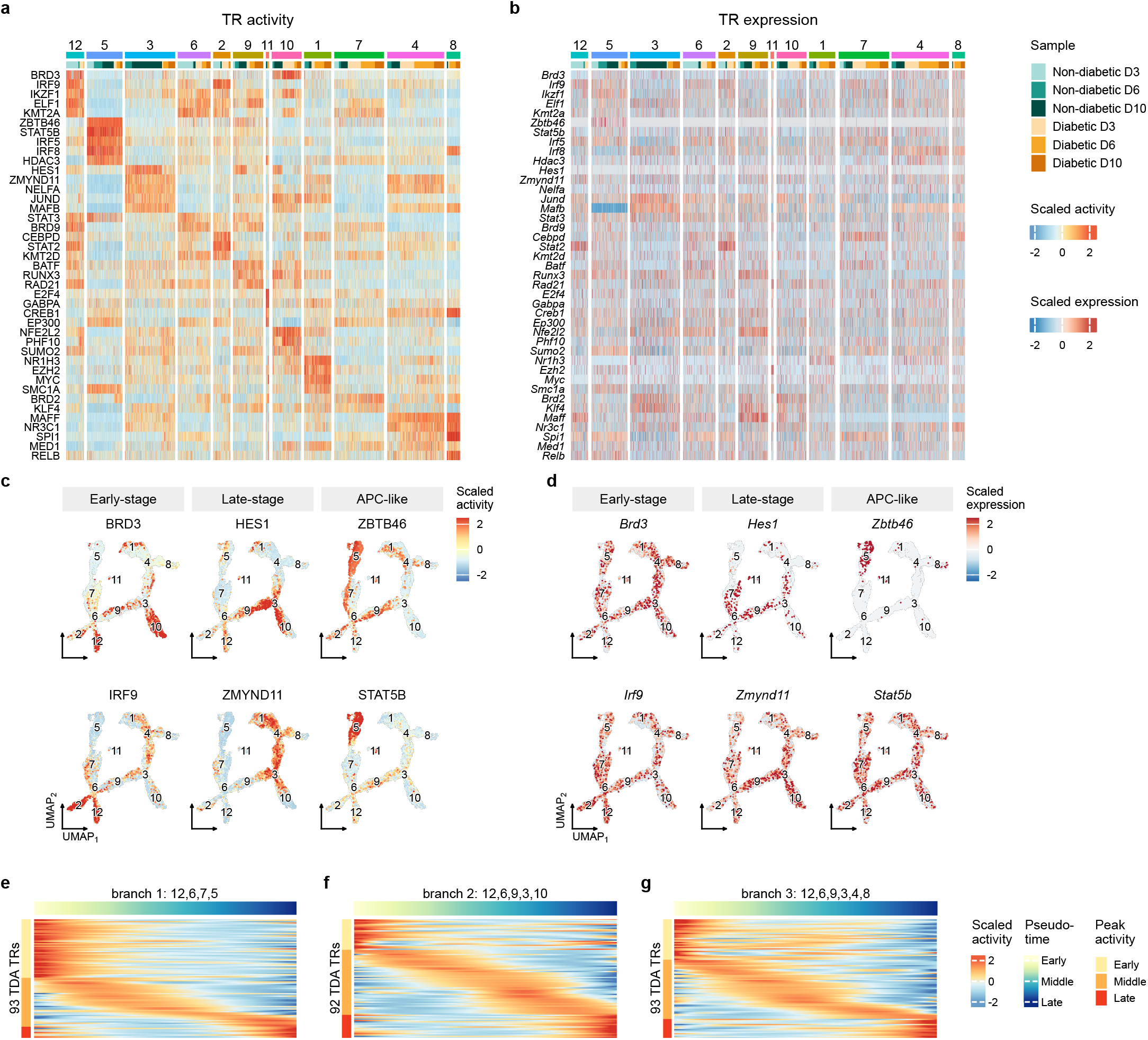
Inferred activity patterns of transcriptional regulators (TRs) differ across wound monocyte/macrophage cell states. **a**. Heatmap visualization of scaled inferred activity of top 5 differentially active TRs per cluster ranked by average log2FC (note that the full list of differentially active TRs can be found in Supplementary Data S3). **b**. Heatmap visualization of scaled expression for respective TR encoding genes. **c**. UMAP visualization of scaled activity of representative differentially active TRs for early-stage, late-stage and APC-like phenotype monocytes/macrophages. **d**. UMAP visualization of scaled expression for respective TR encoding genes.

Cluster 12, which expressed an early-stage interferon activation signature, showed preferential inferred activity of IRF9 and BRD3, which both promote the transcription of inflammatory cytokines, and promote the type I interferon response [35, 36] (Figure 4a-d). Cluster 3, which expressed a late-stage reparative signature, showed preferential inferred activity of HES1 and ZMYND11. HES1 is a transcriptional repressor that downregulates the expression of inflammatory genes [37] and ZMYND11 is a lesser known TF that may negatively regulate the NFkB pathway [38]; thus, both may contribute to the transition of pro-inflammatory to reparative M*φ* phenotypes. Cluster 5, which expressed an APC-like signature, showed preferential inferred activity of ZBTB46 and STAT5B. Interestingly, ZBTB46 is not thought to be expressed in Mo/M*φ*, but is important for maintaining a dendritic cell identity [39]; *Zbtb46* was highly expressed only in cluster 5 cells. STAT5B may promote a tolerogenic state of dendritic cells, leading to enhanced T regulatory cell activity [34].

Cluster 10, which expressed a mixed signature, exhibited preferential inferred activity of NFE2L2 and PHF10 (Fig. 4a, S3). NFE2L2, or nuclear factor erythroid 2–related factor 2, is a master regulator for protection against oxidative stress, but also can downregulate pro-inflammatory cytokines [40, 41]. PHF10 may be important in the development of myeloid progenitors in bone marrow [42] and is part of a chromatin remodeling complex that is important for activation of inflammatory genes [43]. Cluster 1, which expressed a lipid handling, foam cell-like phenotype, showed preferential inferred activity of NR1H3 and EZH2. NR1H3 encodes Liver X Receptor alpha, which is important for lipid metabolism and inflammatory gene regulation [44]. EZH2 is an epigenetic regulator that can repress either pro- or anti-inflammatory genes through histone methylation [45]. Clusters 4, 7, 8, which expressed mixed inflammatory phenotypes, showed preferential inferred activity of BRD2 and NR3C1. BRD2, like its cousin BRD3, plays a key role in chromatin remodeling and promotes expression of inflammatory genes [46] and NR3C1 encodes the glucocorticoid receptor, which is a potent suppressor of inflammatory gene expression [47]. Thus, in these clusters, inferred TR activity predicts mixed and perhaps conflicting pro- and anti-inflammatory functions in DB wound Mo/M*φ* throughout the time course examined. Finally, we identified TR with preferential inferred activity during early, middle and late pseudotimes (Fig. 4e-g, Supplementary Data S6), highlighting additional potential regulators of Mo/M*φ* cell state transitions in wounds.

### In silico TR knockout predicts impact on Mo/M*φ* state transitions

CellOracle was used to predict how knockout of TRs in the gene regulatory network generated for the BITFAM analysis affects Mo/M*φ* state transitions (Fig. 5). First, we calculated perturbation scores (PS) for knockouts of each TR, then summed the negative or positive PS along each trajectory branch [24]. Negative PS scores reflect inhibition of state transitions and positive PS scores reflect enhancement of state transitions along each branch. Figure 5a shows a plot of the negative PS sum for each TR along branch 1 versus 2, which highlights the predicted impact of TR knockouts that inhibit state transitions along both branches (e.g. *Cebpa, Irf8;* Supplementary Video 1), those that have stronger inhibition along branch 1 than branch 2 (e.g. *Irf4, Ogt;* Supplementary Video 2), those that have stronger inhibition along branch 2 than branch 1 (e.g. *Nr1h3, Nr3c1*), and those that have little influence on either branch (e.g. *Kdm1a, Cdk9;* Supplementary Video 3). Figure 5b shows a plot of the positive PS sum for TRs along branch 1 versus 2, which highlights the impact of TR knockouts that strongly promote state transitions along both branches (e.g. *Flk1, Ncor2*), those that have stronger positive influence along branch 1 than branch 2 (e.g. *Cebpb, Atf4*), those that have stronger influence along branch 2 than branch 1 (e.g. *Ogt, Cebpa*), and those that have little influence on either branch (e.g. *Kdm1a, Cdk9*). Comparing branches 2 and 3, knockout of *Cebpa, Irf8* again induced strong inhibition along both branches, whereas knockout of *Fli1* and *Nfatc2* induced stronger inhibition along branch 3 than branch 2, and knockout of *Bhlhe40* and *Nfkb1* induced stronger inhibition along branch 2 than branch 3 (Fig. S4).

**Figure 5.**
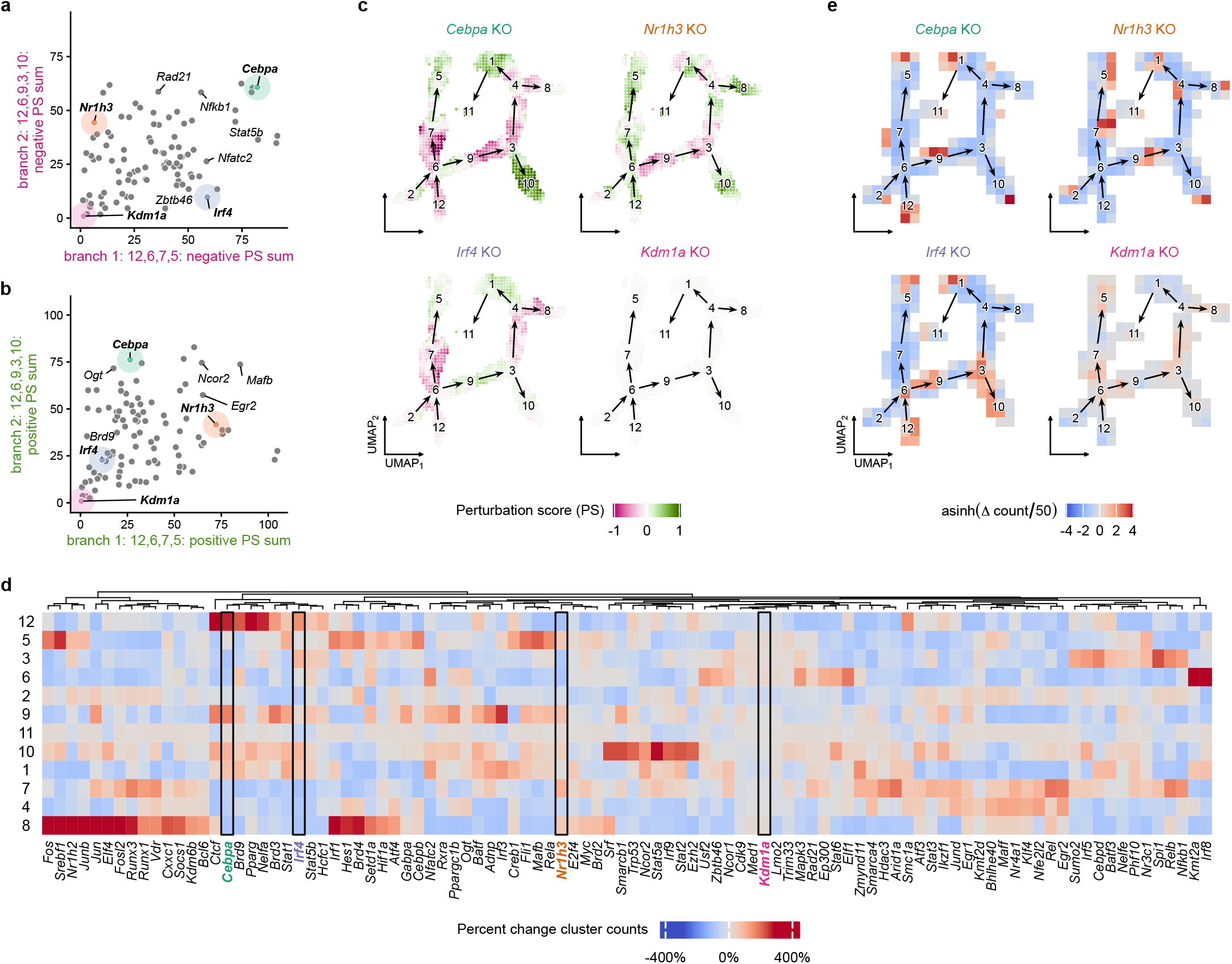
Predicted impact of in silico knockout (KO) of transcriptional regulators (TRs) on altered monocyte/-macrophage cell state transitions. **a**. Scatter plot summarizing predicted relative impact of KO of each of 98 TRs on inhibition of cell state transitions (summed negative perturbation scores (PS)) along branch 1: 12,6,7,5 and branch 2: 12,6,9,3,10. **b**. Scatter plot summarizing predicted relative impact of KO of each of 98 TRs on promotion of cell state transitions (summed positive PS) along branch 1: 12,6,7,5 and branch 2: 12,6,9,3,10. **c**. UMAP visualizations of PS along each trajectory branch for representative TR KOs. Red indicates negative PS (inhibition of state transition), green indicates positive PS (promotion of state transition). **d**. Heatmap visualization showing predicted impact of KO of each of 98 TR on percent change in cluster counts between initial and final states of Markov random walk simulations. Red indicates increase in cell count, blue indicates decrease in cell count. **e**. UMAP visualization of cell density changes of Markov random walk simulations for representative TR KOs.

We note that knockout of many TRs were predicted to inhibit state transitions along portions of each branch, but promote state transitions along other portions. For example, knockout of *Cebpa* showed inhibition of state transitions from early stage through intermediate stages along each trajectory branch, but promoted state transitions within clusters 1 and 10 which were termini of branches 2 and 3 (Fig. 5c). In contrast, other TRs showed uniform inhibition or promotion of state transitions along specific trajectory branches. For example, knockout of *Irf4* showed uniform inhibition of state transitions along trajectory branch 1 (Fig. 5c, Supplementary Video 2). For experimental validation, in a separate study, we showed that knockout of *Irf4* in monocytes using *Csf1r*^Cre^*Irf4* ^fl/fl^ mice reduced differentiation of Mo into cells with the APC phenotype (i.e. along trajectory branch 1 in the current study) [25], consistent with the CellOracle in silico TR knockout prediction.

Next, we performed Markov random walk simulations for each perturbation. These simulations predicted that knockout of selected TRs would result in accumulation of cells in specific clusters (Fig. 5d,e; Supplementary Data S7). For example, individual knockouts of a number of TRs including *Cebpa, Pparg, Ctcf* or *Nelfa* each predicted robust accumulation of cells in the early-stage inflammatory cluster 12 with reductions in other clusters, indicating that knockout of these TRs may result in impaired cell state transitions of early wound Mo/M*φ* to later stage phenotypes. In addition, individual knockouts of *Irf8* or *Kmt2a* predicted robust accumulation of cells in the intermediate cluster 6, indicating that these TRs may be required to transit through this cell state to later stage phenotypes. In contrast, individual knockouts of another set of TRs resulted in accumulation of cells in cluster 5, including *Mafb, Rela* and *Fli1* indicating that these TRs may inhibit cell state transition of Mo into the APC phenotype or may promote differentiation into other phenotypes. Interestingly, individual knockout of a group of TRs including *Spi1, Relb, Nfkb1, Cebpd and Irf5* resulted in accumulation of cells into the reparative late-stage cluster 3, which provides insight into targets for inducing this phenotype to promote tissue repair. In contrast, knockout of *Zmynd11, Ncor2, Nfatc2, Irf3* and *Adnp* induced accumulation of cells in the lipid handling, foam cell-like phenotype cluster 1, populated primarily by late-stage DB cells. Finally, individual knockout of a large group of TRs, including *Fos, Jun, JunB, Srebp1, Nr1h2, Elf4, Fosl2 and Irf1* resulted in accumulation of cells in cluster 8, containing Lyve-1^+^ mixed phenotype cells, another phenotype expressed by late-stage DB cells.

Mo/M*φ* TRs that have been studied in the context of skin wound healing include MAFB, which appears to promote a reparative M*φ* phenotype; experimental knockout of *Mafb* in Mo/M*φ* using *LysM* ^Cre^*Mafb*^fl/fl^ mice led to reduced expression of *Arg1, Il10, Ccl2* and *Ccl12* in IL4/IL13 stimulated bone-marrow derived M*φ* compared to control cells [48]. Additionally, *Mafb* has been shown to be essential for differentiation of human Mo into M*φ*, whereas *Irf4* is essential for Mo differentiation to into Mo-DC [49]. The current Markov random walk analysis (Fig. 5d) predicted that *Mafb* knockout would result in increased accumulation of cells in cluster 5 (i.e. the APC or Mo-DC phenotype; Supplementary Video 4), whereas *Irf4* knockout promotes accumulation of cells in cluster 3 (i.e. the reparative phenotype), suggesting that these TRs promote alternative state transitions from early to late wound stage phenotypes. In addition, macrophage-specific knockout of *Pparg* resulted in prolonged wound inflammation and impaired healing [50]. The Markov random walk analysis predicted that *Pparg* knockout would result in accumulation of pro-inflammatory cluster 12 cells, consistent with the experimental results.

### In silico TR overexpression predicts impact on Mo/M*φ* state transitions

CellOracle was used also to simulate how TR overexpression may affect Mo/M*φ* state transitions (Fig. 6). In overexpression simulations, the expression of the gene encoding the TR is universally set to an upper limit derived from the observed single-cell expression range of that gene. In general, overexpression of TRs was predicted to induce inverse effects compared to TR knockout on Mo/M*φ* state transitions. For TR knockouts predicted to inhibit state transitions along both trajectory branches, overexpression was predicted to promote state transitions, albeit to different degrees (Fig. 6a; e.g. *Cebpa, Irf8*). For knockouts predicted to have stronger influence along branch 1 than branch 2 (e.g. *Irf4, Ogt*), and those that have stronger influence along branch 2 than branch 1 (e.g. *Nr1h3, Nr3c1*), overexpression of these TRs promoted state transitions along these branches. Conversely, for TR knockouts predicted to promote state transitions along both trajectory branches, overexpression was predicted to inhibit these state transitions (Fig. 6b; e.g. *Flk1, Ncor2*). For knockouts predicted to have stronger influence on branch 1 than on branch 2 (e.g. *Cebpb, Atf4*), and those predicted to have stronger influence on branch 2 than on branch 1 (e.g. *Ogt, Cebpa*), overexpression of these TRs inhibited state transitions along these branches.

**Figure 6.**
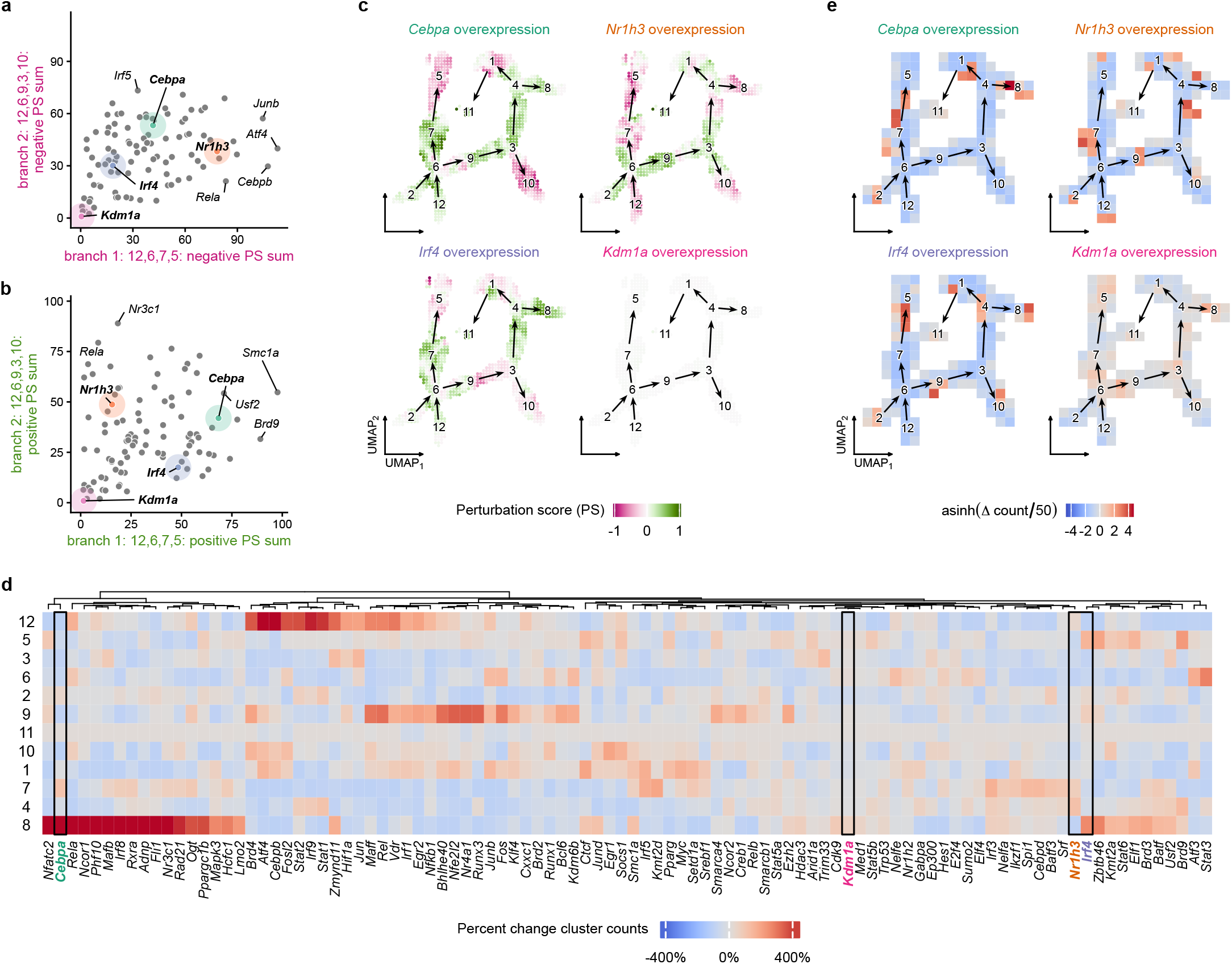
Predicted impact of in silico overexpression (OE) of transcriptional regulators (TRs) on altered monocyte/macrophage cell state transitions. **a**. Scatter plot summarizing predicted relative impact of OE of each of 98 TRs on inhibition of cell state transitions (summed negative perturbation scores (PS)) along branch 1: 12,6,7,5 and branch 2: 12,6,9,3,10. **b**. Scatter plot summarizing predicted relative impact of OE of each of 98 TRs on promotion of cell state transitions (summed positive PS) along branch 1: 12,6,7,5 and branch 2: 12,6,9,3,10. **c**. UMAP visualizations of PS along each trajectory branch for representative TR OEs. Red indicates negative PS (inhibition of state transition), green indicates positive PS (promotion of state transition). **d**. Heatmap visualization showing predicted impact of OE of each of 98 TR on percent change in cluster counts between initial and final states of Markov random walk simulations. Red indicates increase in cell count, blue indicates decrease in cell count. **e**. UMAP visualization of cell density changes of Markov random walk simulations for representative TR OEs.

Markov random walk simulations predicted that overexpression of selected TRs would result in accumulation of cells in specific clusters (Fig. 6c; Supplementary Data S8). Overexpression of members of the Interferon-Stimulated Gene Factor 3 complex (*Stat1, Stat2* and *Irf9*), along with *Nfkb1, Cebpb, Fosl2*, all increased cells in early pro-inflammatory cluster 12. Overexpression of *Irf4, Zbtb46* and *Brd9* increased cells with an APC phenotype in cluster 5, and overexpression of *Trim33, Zmynd11* and *Jun* produced modest increases in cells with a reparative phenotype in cluster 3. Finally, overexpression of *Irf5, Ctcf* and *Atf4* produced an increase in cells with a lipid handling, foam cell-like phenotype, and overexpression of a large selection of TRs (e.g. *Cebpa, Nfatc2, Rela, Irf8, Mafb*) each resulted in an increase in cells in the Lyve-1 phenotype cluster 8.

## 4 Discussion

Although dysregulation of Mo/M*φ* activity is known to contribute to impaired healing in diabetes [11, 12], the mechanisms underlying this dysregulation are not well understood. In the present study, we used the Lamian computational framework [23] and our newly developed Pseudotime Graph Diffusion method on our scRNA-seq data to comprehensively identify alterations in Mo/M*φ* phenotypes over the course of healing, and show that state transitions from early stage inflammatory phenotypes to later stage reparative and APC phenotypes are impaired and that transitions to inflammatory, foam cell-like, and Lyve-1^+^ phenotypes are enhanced in wounds of diabetic mice. Using our BITFAM model [21, 22], we also identified a broad range of TRs predicted to be preferentially active in each cell state. We used CellOracle [24] to perform in silico TR perturbation to predict which TRs drive cell state transitions in non-diabetic and diabetic wounds; certain TRs were predicted to drive cell state transitions along multiple trajectories (e.g. CEBPA, IRF8), whereas other TRs were predicted to drive cell state transition towards reparative phenotypes (e.g. NR1H3, NR3C1) and or towards an antigen-presenting phenotype (e.g. IRF4, OGT). Finally, we validated selected findings using existing experimental data, confirming the usefulness of this approach for identifying TRs that drive state transitions in both non-diabetic and diabetic wounds.

Other groups have recently published findings on the heterogeneity of wound Mo/M*φ* in both diabetic mice and humans. Using the streptozotocin model of type 1 diabetes and scRNA-seq [51], six Mo/M*φ* subsets and two dendritic cell subsets were observed in skin swounds, with varying contributions of ND and DB cells to each cluster over the time points observed (one to seven days post-five mm excisional wounding). Although a list of TFs that were expressed in each cluster was provided, TF activity was not assessed, and findings from the present study indicate that TF expression levels do not correlate well with predicted TF activity. Another study profiled human wound cells during normal skin wound healing (one to 30 days post four mm excisional wounding) [52], and found that, similar to what occurs in rodent models, Mo/M*φ* progress from pro-inflammatory to pro-resolution phenotypes as healing progresses; however the role of TFs in this switch was not a focus of the study. Another study profiled wound cells from human diabetic foot ulcers and found that Mo/M*φ* classified as “M1” were underrepresented and those classified as “M2” were overrepresented in wounds that did not heal compared to those that healed [32], highlighting dysregulation of Mo/M*φ* in non-healing wounds. This study was limited by the oversimplified and biased categorization of wound Mo/M*φ*, and the role of TFs in driving these phenotypes was not investigated. Our study adds to the published literature by comprehensively assessing the TRs that drive the heterogeneous phenotypes of wound Mo/M*φ* in both ND and DB wounds.

A number of recent studies have demonstrated that epigenetic gene regulation via histone modifications, DNA modifications, and microRNA can influence macrophage state transitions during wound healing [12, 53]. Because epigenetic regulators are included in the ChIP-Atlas database, these were present in both our BITFAM and CellOracle analyses, and our findings highlight the importance of these regulators in Mo/M*φ* state transitions during wound healing. For example, the histone methylase KMT2A, or MLL1, adds methyl groups to histone H3 at K4, thereby promoting chromatin opening and subsequent gene expression. We found that KMT2A was predicted to be highly active in the early-stage pro-inflammatory cluster 12 and to a lesser extent in intermediate-stage clusters 6 and 7, and that in silico knockout of *Kmt2a* was predicted to reduce cells in cluster 12, as well as terminal clusters 3 and 5, and increase cells in cluster 6 and to a lesser extent cluster 1, the latter expressing the DB-associated lipid handling phenotype that was less inflammatory than cluster 12. Consistent with these findings, Mo/M*φ* specific *Mll1* knockout mice exhibited reduced wound macrophage inflammatory cytokine production and delayed wound healing [54].

Additional epigenetic regulators predicted to be important in specifying Mo/M*φ* states, which have been less well-studied during wound healing, include Bromodomain-containing protein (BRD) proteins, which are epigenetic readers considered to be master regulators of gene expression, O-GlcNAc transferase (OGT), which adds O-GlcNAc to TFs and other proteins, generally suppressing pro-inflammatory responses, and Zinc Finger MYND-Type Containing 11 (ZMYND11), which acts as a transcriptional corepressor of inflammatory genes that are marked with the histone modification H3.3K36me3. These results highlight the wide variety of factors that contribute to specifying wound Mo/M*φ*, many of which have not been studied in the context of impaired wound healing with diabetes.

A limitation of our study is that, although our results provide unbiased and comprehensive insight into the regulation of Mo/M*φ* cell states in normal and impaired wound healing, how the TRs identified as being important in specifying Mo/M*φ* cell states are themselves regulated awaits further study. Another limitation of our study includes inability to differentiate between resident and infiltrating cells that contribute to the wound Mo/M*φ* pool. The question of whether resident and infiltrating Mo/M*φ* exhibit different phenotypes in ND and DB wounds, and whether different regulatory mechanisms are involved in cells from different sources will be a topic of future study. Other limitations of this study relate to the computational analyses. First, our trajectory inference relied on a single method (Lamian), which constructs a minimum spanning tree across both ND and DB cells jointly. This tree-based topology assumes linear progression from an initial cluster to a limited number of terminal clusters, precluding more complex dynamics such as phenotypic reversion or cyclical state transitions. Moreover, although every cluster contains cells from both conditions, the joint construction may suggest transitions that do not occur within either single condition. The trajectory modeled here therefore represents possible, but not exhaustive, cell state transitions agnostic to condition. In addition, our CellOracle perturbation analyses lack formal statistical testing; perturbation scores should be interpreted as computational predictions that prioritize candidates for experimental validation rather than as statistically confirmed effects.

In conclusion, we identified TRs that likely promote or inhibit Mo/M*φ* state transitions towards desirable and undesirable phenotypes in ND and DB wounds. These findings provide insight into novel targets for altering Mo/M*φ* phenotypes to promote healing of DB wounds.

## Acknowledgements

This study was supported by the NIGMS through grant R35GM136228 to TJK, and the NIDDK Diabetic Complications Consortium through grants DK076169 and DK115255 to YD and TJK.

## Author Contributions

BL and JP contributed equally to this work. BL developed bioinformatics framework, implemented workflow, analyzed data, and wrote manuscript. JP performed experiments, analyzed data, and wrote manuscript. YD designed study, developed bioinformatics framework, analyzed data, and wrote manuscript. TJK designed study, analyzed data, and wrote manuscript.

## SUPPLEMENTARY INFORMATION

**Supplementary Figure 1. Pathway enrichment analysis of temporally dynamic genes across pseudotime branches**.

**Supplementary Figure 2. Feature plots of canonical marker genes across monocyte/-macrophage clusters**.

**Supplementary Figure 3. Feature plots of marker transcriptional regulators with differentially inferred activity**.

**Supplementary Figure 4. CellOracle perturbation analysis of transcriptional regulators across pseudotime branches**.

**Supplementary Data 1. Cluster-based differential gene expression analysis. Results of differential expression testing across monocyte/macrophage clusters**.

**Supplementary Data 2. Cluster-based gene set enrichment analysis (GSEA). Supplementary Data 3. Temporally dynamic genes along pseudotime branches**.

**Supplementary Data 4. Cluster-based differential transcriptional regulator (TR) activity**.

**Supplementary Data 5. Cluster-level pseudobulk transcriptional regulator (TR) expression**.

**Supplementary Data 6. Temporally dynamic transcriptional regulators (TRs) along pseudotime branches**.

**Supplementary Data 7. Markov chain transition changes under transcriptional regulator (TR) knockout**.

**Supplementary Data 8. Markov chain transition changes under transcriptional regulator (TR) overexpression**.

**Supplementary Video 1. Simulated particle flow along pseudotime and simulated Cebpa knockout vector fields**.

**Supplementary Video 2. Simulated particle flow along pseudotime and simulated Irf4 knockout vector fields**.

**Supplementary Video 3. Simulated particle flow along pseudotime and simulated Kdm1a knockout vector fields**.

**Supplementary Video 4. Simulated particle flow along pseudotime and simulated Mafb knockout vector fields**.

