## Supplementary Information for "Transcriptional regulators predicted to drive macrophage dysregulation during impaired wound healing in diabetic mice"

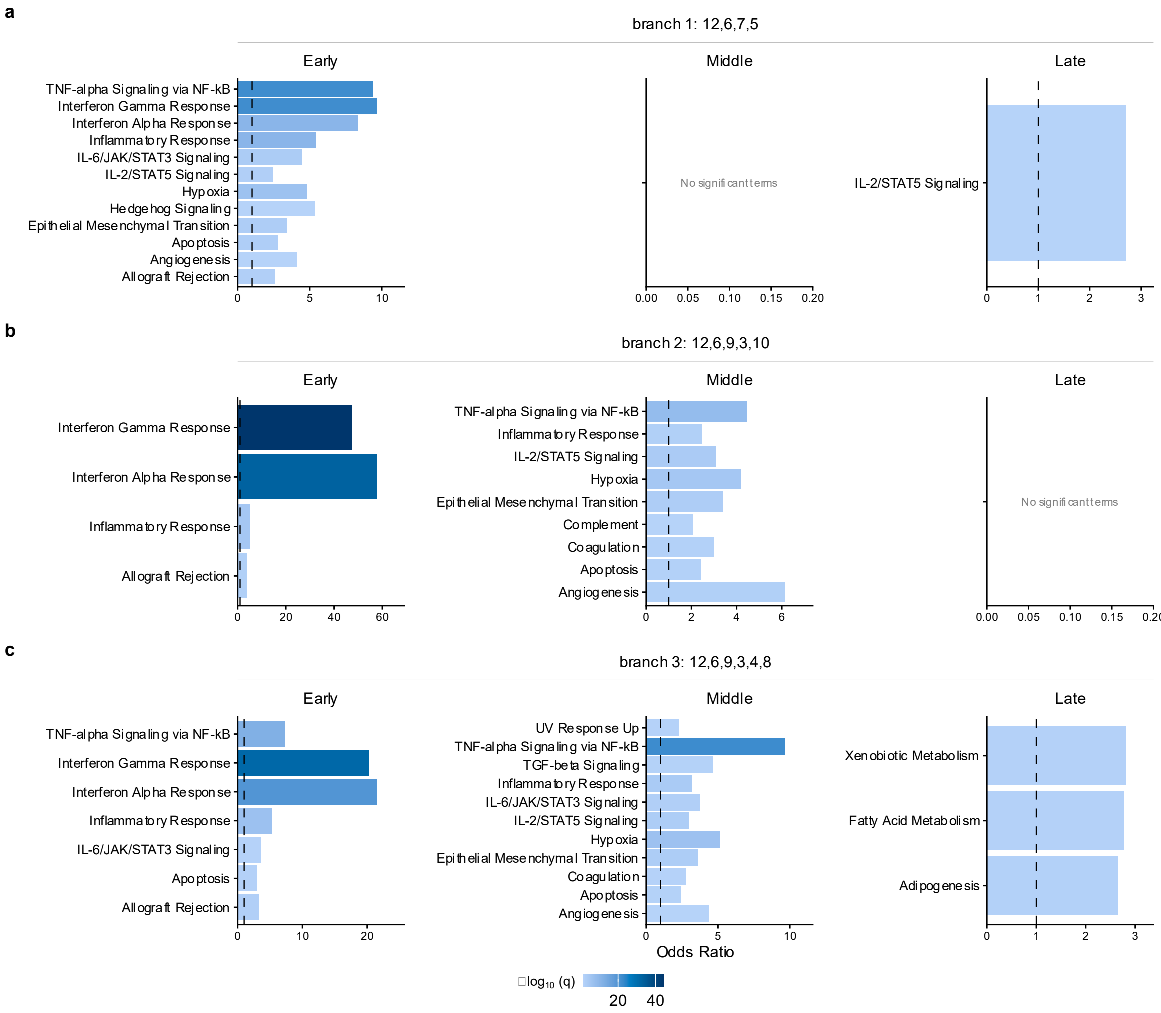


**Supplementary Figure 1. Pathway enrichment analysis of temporally dynamic genes across pseudotime branches.** For each branch, Lamian temporally dynamic genes (TDEs; FDR < 0.05) were classified by the pseudotime location of their peak expression into early, middle, or late groups. Overrepresentation analysis was performed against the MSigDB Hallmark gene set collection. Bar length indicates odds ratio; color indicates statistical significance (-log10 q-value). All significant terms ($q < 0.05$) are shown. (a) Branch 1 (clusters 12, 6, 7, 5). (b) Branch 2 (clusters 12, 6, 9, 3, 10). (c) Branch 3 (clusters 12, 6, 9, 3, 4, 8).

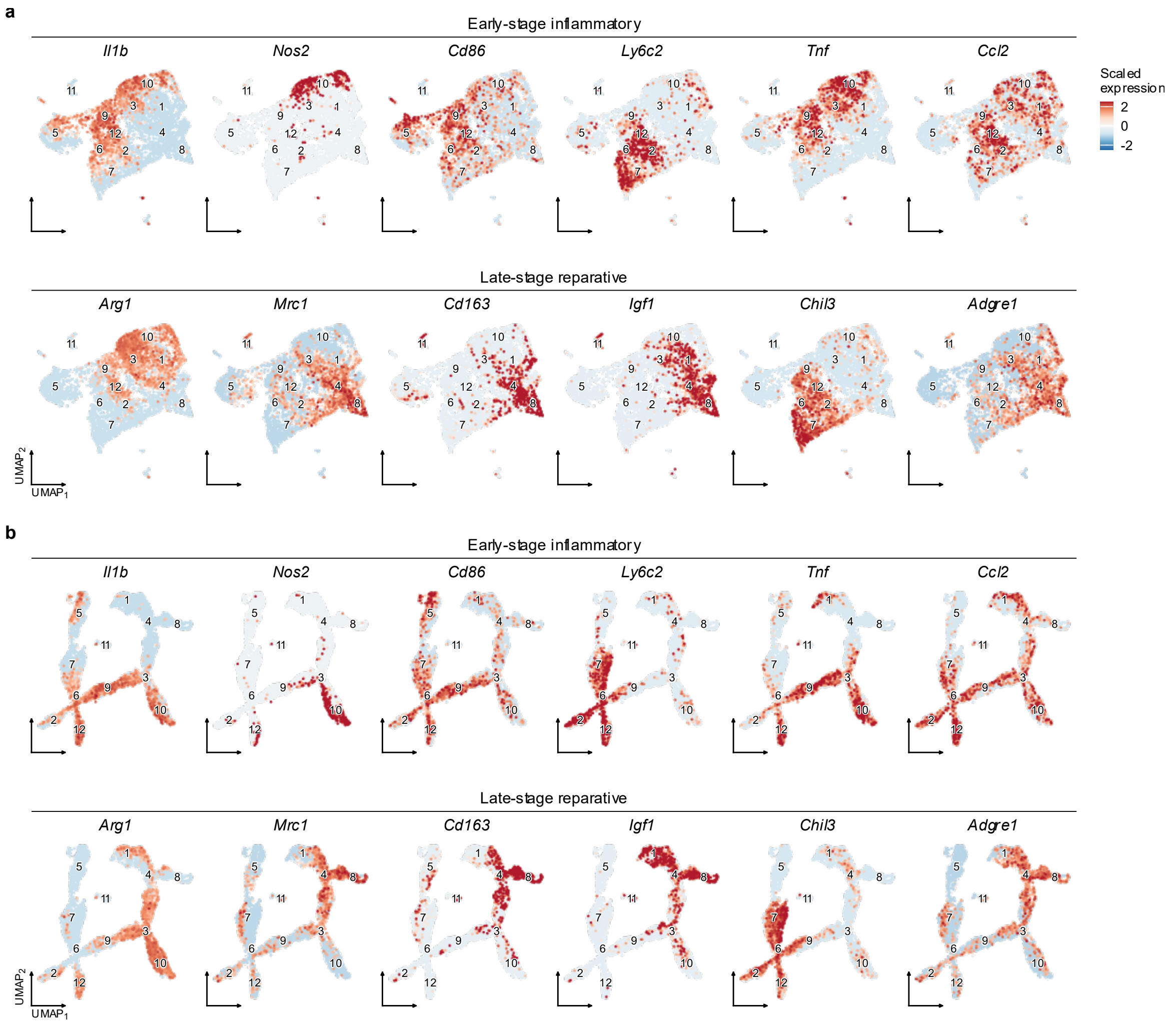


**Supplementary Figure 2. Feature plots of canonical marker genes across monocyte/macrophage clusters.** (a) UMAP embedding with standard layout showing scaled expression of selected early-stage inflammatory (*Il1b*, *Nos2*, *Cd86*, *Ly6c2*, *Tnf*, *Ccl2*) and late-stage reparative (*Arg1*, *Mrc1*, *Cd163*, *Igf1*, *Chil3*, *Adgre1*) marker genes. (b) UMAP embedding after Pseudotime Graph Diffusion (PGD)-smoothing, preserving local neighborhood structure while enhancing trajectory continuity. Color scale indicates scaled expression level.

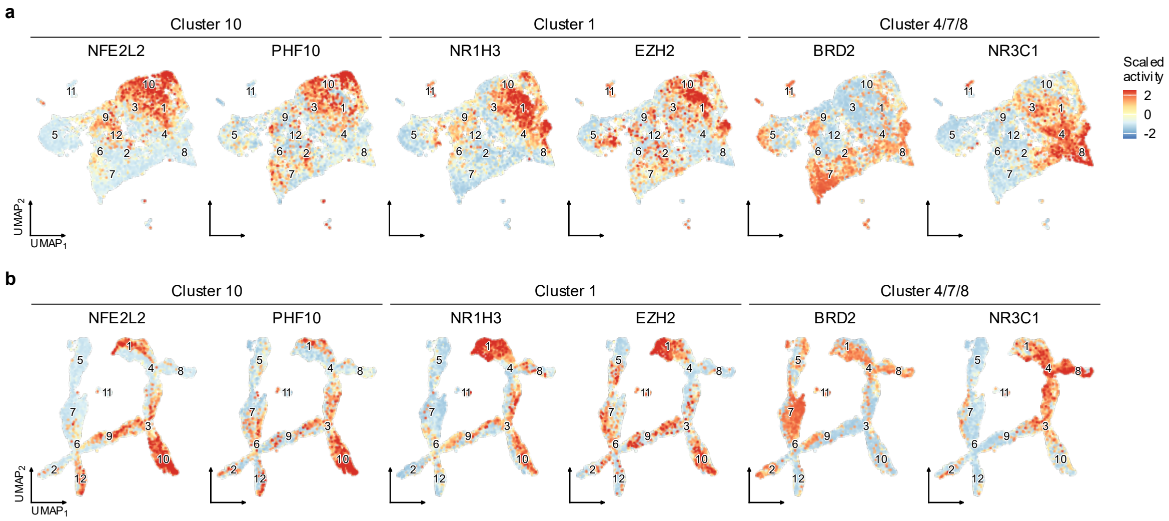


**Supplementary Figure 3. Feature plots of marker transcriptional regulators with differentially inferred activity.** Transcriptional regulator (TR) activity was inferred using BITFAM, and median activity per TR per cell across four independent runs is shown. Marker TRs were identified as differentially active in specific clusters relative to all other clusters. (a) Standard UMAP layout. (b) UMAP layout after Pseudotime Graph Diffusion (PGD)-smoothing. TRs are grouped by their marker cluster: NFE2L2 and PHF10 (cluster 10), NR1H3 and EZH2 (cluster 1), BRD2 and NR3C1 (clusters 4/7/8). Color scale indicates scaled activity level.

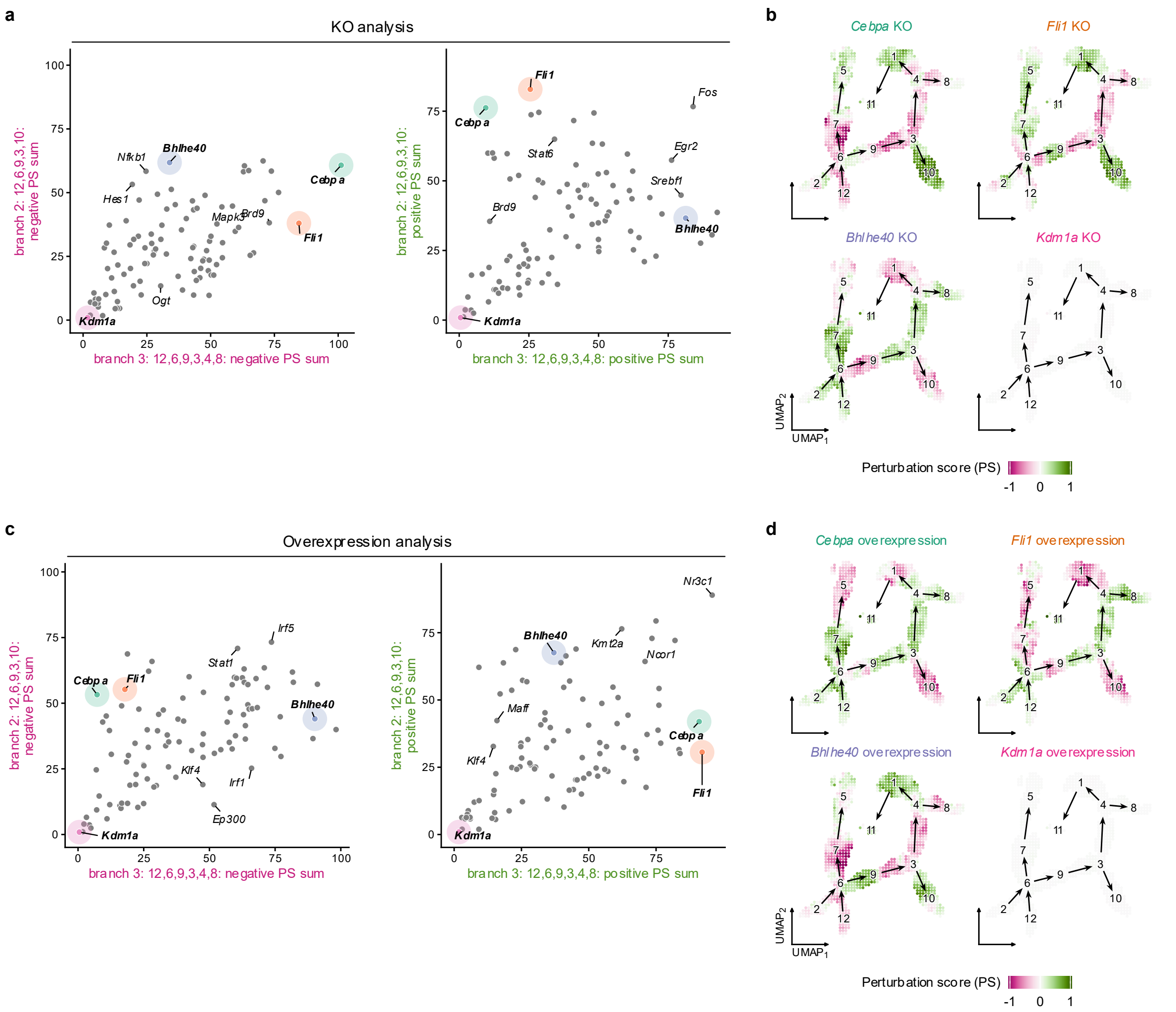


**Supplementary Figure 4. CellOracle perturbation analysis of transcriptional regulators across pseudotime branches.** Systematic in silico knockout (KO) and overexpression of 98 TRs was performed using CellOracle. Perturbation scores (PS) were computed along each branch and summarized as negative PS sum (inhibition of cell state transitions) and positive PS sum (promotion of cell state transitions). (a) Scatter plots of summarized negative (left) and positive (right) PS sums for each TR under KO, comparing branch 2 (y-axis) and branch 3 (x-axis). Select TRs are labeled. (b) PGD-smoothed UMAP projections of grid-level PS values for four representative TRs (*Cebpa*, *Fli1*, *Bhlhe40*, *Kdm1a*) under KO, illustrating the spatial distribution of perturbation effects along branches. (c) As in (a), but for overexpression perturbations. (d) As in (b), but for overexpression of the same four representative TRs.

**Supplementary Data 1. Cluster-based differential gene expression analysis.** Results of differential gene expression testing across cell clusters. Sheet 1 (“one_vs_rest”): one-vs-rest marker gene analysis for each cluster. Sheet 2 (“db_vs_nd”): diabetic vs. non-diabetic differential expression within each cluster. Sheet 3 (“combined”): merged results from both analyses. Columns include log fold-change, p-values, and adjusted p-values.

**Supplementary Data 2. Cluster-based gene set enrichment analysis (GSEA).** GSEA results for each cluster, including enrichment scores, normalized enrichment scores (NES), p-values, and adjusted p-values.

**Supplementary Data 3. Temporally dynamic genes along pseudotime branches.** Lamian trajectory differential expression (TDE) analysis results. Sheet 1 (“TDE”): temporally dynamic genes (FDR < 0.05) identified along each pseudotime branch. Sheet 2 (“TDE_Pathway_Enrichment”): Gene Ontology overrepresentation analysis of TDE genes.

**Supplementary Data 4. Cluster-based differential transcriptional regulator (TR) activity.** Statistical significance of differential TR activity across clusters, based on BITFAM-inferred activity scores aggregated across four independent runs.

**Supplementary Data 5. Cluster-level pseudobulk transcriptional regulator (TR) expression.** Log-normalized pseudobulk RNA expression of TRs aggregated by cluster.

**Supplementary Data 6. Temporally dynamic transcriptional regulators (TRs) along pseudotime branches.** Lamian TDE analysis of BITFAM-inferred TR activity scores along each pseudotime branch. Includes peak pseudotime position and temporal classification (early, middle, late) for each TR per branch.

**Supplementary Data 7. Markov chain transition changes for transcriptional regulator (TR) knockout.** Percent change in Markov chain cluster occupancy after 500 simulation steps following in silico CellOracle knockout of each TR, relative to baseline (step 0).

**Supplementary Data 8. Markov chain transition changes for transcriptional regulator (TR) overexpression.** Percent change in Markov chain cluster occupancy after 500 simulation steps following in silico CellOracle overexpression of each TR, relative to baseline (step 0).

---

**Supplementary Video 1. Simulated particle flow indicating cell state transitions along pseudotime and simulated *Cebpa* knockout vector fields.** Left: Note particle flow indicating cell state transitions from early stage cells (lower left) to APC phenotype cells (upper left) and reparative, lipid handling and LYVE1+ phenotypes (right). Right: Note reversal of particle flow (marked by red) at intermediate stages between early stage cells and APC phenotype cells, and between early stage cells and other late stage phenotype cells, indicating inhibition of cell state transitions predicted to be induced by *Cebpa* knockout. Particle flow is enhanced (marked by green) at other parts of the cell state trajectory.

**Supplementary Video 2. Simulated particle flow indicating cell state transitions along pseudotime and simulated *Irf4* knockout vector fields.** Left: Note particle flow indicating cell state transitions from early stage cells (lower left) to APC phenotype cells (upper left) and reparative, lipid handling and LYVE1+ phenotypes (right). Right: Note reversal of particle flow (marked by red) at intermediate stages between early stage cells and APC phenotype cells, indicating inhibition of cell state transition predicted to be induced by *Irf4* knockout. Particle flow is enhanced (marked by green) at other parts of the cell state trajectory.

**Supplementary Video 3. Simulated particle flow indicating cell state transitions along pseudotime and simulated *Kdm1a* knockout vector fields.** Left: Note particle flow indicating cell state transitions from early stage cells (lower left) to APC phenotype cells (upper left) and reparative, lipid handling and LYVE1+ phenotypes (right). Right: Note little predicted impact of *Kdm1a* knockout on particle flow (white), indicating little impact on cell state transitions.

**Supplementary Video 4. Simulated particle flow indicating cell state transitions along pseudotime and simulated *Mafb* knockout vector fields.** Left: Note particle flow indicating cell state transitions from early stage cells (lower left) to APC phenotype cells (upper left) and reparative, lipid handling and LYVE1+ phenotypes (right). Right: Note reversal of particle flow (marked by red) at intermediate stages between early stage cells and reparative phenotype cells, indicating inhibition of cell state transition predicted to be induced by *Mafb* knockout. Particle flow is enhanced (marked by green) at other parts of the cell state trajectory, particularly towards APC phenotype cells.
